# Genomic Epidemiology of Dengue Virus in Ivory Coast Reveals Five Decades of Viral Circulation with Repeated Introductions from Burkina Faso

**DOI:** 10.64898/2026.11.18.767690

**Authors:** Xinghua Tao, Herve A. Kadjo, Sylla Yahaya, Kamo Emily, Adjogoua Edgard, Marc A. Suchard, Nidia S. Trovao

## Abstract

Dengue fever is a rapidly emerging mosquito-borne disease in Africa, yet the continent remains critically underrepresented in genomic surveillance efforts. This disparity is especially pronounced in West Africa, despite phylogenetic studies being concentrated in only a few countries, such as Burkina Faso. Within this broader regional gap, researchers have yet to systematically characterize the dengue virus (DENV) lineages circulating in the Ivory Coast. In this study, we present 24 newly sequenced DENV genomes from Ivory Coast, spanning serotypes DENV-1, DENV-2, and DENV-3, collected between 2017 and 2023. Combined with publicly available sequences, we assembled the most comprehensive dataset to date for the country.

We performed maximum-likelihood and Bayesian phylogeographic analyses using global background datasets and reconstructed the spatial and temporal dynamics of dengue virus circulation in Ivory Coast. Our findings reveal multiple independent introductions of DENV into Ivory Coast, predominantly from neighboring Burkina Faso. We identify five introductions of DENV-1 between 2017 and 2019 and four introductions of DENV-2 between 2015 and 2016. DENV-3 showed two separate introductions, one in 2015 and another in 2018. Notably, we also identified an early DENV-1 introduction from India around 1977, suggesting long-distance historical viral movement likely linked to colonial and post-colonial military presence. These results underscore the need for sustained genomic surveillance to monitor viral evolution and inform regional public health responses aimed at controlling dengue outbreaks in West Africa.

## 1 Background

Dengue fever is a mosquito-borne viral illness that poses a major public health concern across tropical and subtropical regions. It is caused by infection with one of four antigenically distinct dengue virus (DENV) serotypes: DENV-1, DENV-2, DENV-3, and DENV-4. Infection with a single serotype induces long-term immunity to that serotype, but only short-lived cross-protection against heterologous serotypes [Guzman and Harris, 2015]. Subsequent infection with a heterologous serotype is associated with an increased risk of severe dengue, including dengue hemorrhagic fever and dengue shock syndrome, with antibody-dependent enhancement (ADE) considered an important contributing mechanism [Guzman et al., 2013]. As a result, monitoring the circulation of multiple serotypes is essential for effective disease surveillance and outbreak preparedness.

Although dengue has been reported in Africa since the 19th century [Diallo et al., 2003], the true burden of disease remains largely underestimated. Historical reports described dengue epidemics in Zanzibar in 1823 and 1870, Egypt in 1887 and 1927, Burkina Faso in 1925, South Africa in 1926–1927, and Senegal in 1927–1928 [Gubler and Clark, 1995, World Health Organization, 2009, Cornet, 1993]. However, except for the South African outbreak, these early epidemics were not laboratory confirmed and therefore remain uncertain [Amarasinghe et al., 2011]. Between 1960 and 2010, only 20 laboratory-confirmed dengue outbreaks were reported across 15 African countries, although nearly 300,000 cases were recorded in several large epidemics, including those in the Seychelles, Reunion, Djibouti, Comoros, and Cape Verde [Amarasinghe et al., 2011]. DENV was first isolated from human patients in Africa in Nigeria during the 1960s [Carey et al., 1971], and all four serotypes have subsequently been detected on the continent, with DENV-2 and DENV-1 responsible for many reported epidemics [Amarasinghe et al., 2011]. Despite this long history of documented transmission, modeling and seroepidemiological studies suggest that dengue circulation in Africa is substantially underestimated [Bhatt et al., 2013, Zingue et al., 2019]. Limited awareness, diagnostic capacity, and misclassification as other febrile illnesses likely contribute to this under-recognition [Dieng et al., 2022b]. Consistent with these surveillance gaps, Africa remains markedly underrepresented in genomic datasets, accounting for less than 1% of globally available DENV sequences [Selhorst et al., 2023], despite being one of the most affected regions [Sharp, 2015].

A recent study covering 2013–2023 found that over 80% of confirmed dengue cases in Africa originated from West Africa, particularly Burkina Faso [Mwanyika et al., 2025]. Most phylogenetic studies to date have focused on countries such as Burkina Faso [Letizia et al., 2022], Kenya [Nyathi et al., 2024, Muthanje et al., 2022], Senegal [Dieng et al., 2022a, Gaye et al., 2021], and Tanzania [Kelly et al., 2023], likely due to the greater availability of genetic sequences from these nations.

In contrast, dengue in Ivory Coast has received comparatively little attention. The earliest virologic evidence of dengue in Ivory Coast dates to 1980, when sylvatic DENV-2 was isolated from wild Aedes mosquitoes; no human infections were detected, and the virus was thought to be maintained in wilderness areas [Roche et al., 1983]. The earliest report documented the isolation of DENV-1 from a soldier who had arrived in Abidjan from France in 1998 [Durand et al., 2000]. During the 2017 outbreak, DENV-2 was identified in travelers returning from Ivory Coast to Japan and Senegal, with phylogenetic analyses indicating that these viruses were closely related to strains circulating during the 2016 dengue outbreak in Burkina Faso [Suzuki et al., 2017, Dieng et al., 2022b]. The substantial disease burden in Ivory Coast has been historically underrecognized in genomic research. Although the first documented case dates back to 1998 [Durand et al., 2000], no comprehensive phylodynamic analysis has been conducted until now.

## 2 Objective

In this study, we aim to characterize the genetic diversity, evolutionary history, and geographic transmission patterns of DENV circulating in Ivory Coast. In this study, we report 24 newly sequenced DENV genomes from Ivory Coast, representing serotypes DENV-1, DENV-2, and DENV-3. By integrating these with 7 existing sequences from GISAID (Global Initiative on Sharing All Influenza Data)[Khare et al., 2021], we assemble the most comprehensive dataset to date for the country. Along with a comprehensive background dataset, this analysis provides novel insights into the genetic diversity and transmission patterns of DENV in Ivory Coast and strengthens the foundation for regional outbreak response and disease control strategies.

## 3 Materials and methods

### 3.1 Sample collection and sequencing

#### 3.1.1 Sample collection

We collect blood samples from febrile patients who present to the outpatient departments of health facilities across 83 health districts nationwide. We identify suspected dengue cases according to the World Health Organization (WHO) case definition. We draw venous blood into plain (red-top, additive-free) tubes. We accompany each sample with a standardized case investigation form that records the patient’s demographic, epidemiological, clinical, and biological data, including the date of symptom onset. We transport samples under cold-chain conditions (2 *−* 8*^◦^*C) using triple-layer packaging, in accordance with WHO guidance on the transport of infectious substances, to the National Reference Laboratory (NRL) for Viral Haemorrhagic Fevers, Department of Epidemic Viruses, Institut Pasteur de Ĉote d’Ivoire. Upon receipt, laboratory personnel centrifuge the samples at 3000 revolutions per minute for 5 minutes and divide the serum into three aliquots of 300 microliters. The first aliquot is transferred to the institutional biobank for long-term storage in liquid nitrogen (*−*196*^◦^*C). The second is stored at *−*70*^◦^*C in a dedicated freezer as a back-up for confirmatory or further testing. The third is used for DENV diagnosis: for samples that we collect within 7 days of symptom onset, we test for DENV RNA by reverse transcription polymerase chain reaction (RT-PCR) using the Trioplex real-time RT-PCR Assay Primer and Probe Set (CDC; catalog #KT0166), which the CDC develops; for samples that we collect later, we screen for DENV-specific IgM antibodies using an IgM antibody-capture enzyme-linked immunosorbent assay (MAC-ELISA). We perform all assays according to the manufacturers’ instructions.

#### 3.1.2 Whole-genome sequencing of dengue virus

We ship all RT-PCR-positive samples on dry ice using triple-layer packaging, in accordance with WHO guidance on the transport of infectious substances [World Health Organization, 2021], to Institut Pasteur de Dakar, Senegal, where we perform whole-genome sequencing using a target-enrichment approach based on hybridization capture with the Twist Comprehensive Viral Research Panel (Twist Bioscience, South San Francisco, CA, USA).

We extract total viral RNA from 200 *µ*L of serum using the QIAamp Viral RNA Mini Kit (Qiagen, Hilden, Germany) and elute it in 60 *µ*L of AVE buffer. We process only samples with a real-time RT-PCR cycle threshold (Ct) not greater than 30. We treat the extracts with DNase I to remove host genomic DNA and quantify the RNA using the Qubit RNA HS Assay Kit (Thermo Fisher Scientific, Waltham, MA, USA). We prepare libraries with the Twist Library Preparation Kit for Viral RNA through random-primed first- and second-strand cDNA synthesis, enzymatic fragmentation, end repair and dA-tailing, and ligation of Twist Universal Adapters with unique dual indexes (UDI), which enable unambiguous sample identification and correction of index hopping. We amplify indexed libraries by PCR (10 cycles), purify them with Twist DNA Purification Beads, and assess their yield using Qubit dsDNA HS and fragment size distribution using TapeStation 4150 (Agilent Technologies, Santa Clara, CA, USA), with an expected peak at 300 *−* 400 bp.

We enrich viral targets through solution-based hybridization capture as described above. This sequence-agnostic approach allows the capture of highly divergent strains and limits the coverage dropout associated with primer mismatches in amplicon-based protocols [Grubaugh et al., 2019]. We pool libraries in equimolar amounts (eight libraries per pool; 1.5 *µ*g total, 187.5 ng per library) and hybridize them to the probes at 70*^◦^*C for 16 h in the presence of Twist universal blockers and adapter blockers. We capture probe-target hybrids on streptavidin beads, perform washes under increasing stringency, and amplify the captured material through 12 *−* 14 cycles of post-capture PCR. We requantify the enriched libraries, check their quality, and normalize them. We perform sequencing on an Illumina MiSeq platform using 2 *×* 150 bp paired-end reads, with a target depth of 1 *−* 2 million reads per sample. We include one extraction blank and one library blank in each run to monitor cross-contamination. We check the quality of raw reads with FastQC [Andrews, 2010], trim adapters and low-quality bases using fastp / Trimmomatic (*Q ≥* 30), and remove human reads in silico by mapping against the GRCh38 human reference genome [Chen et al., 2018, Bolger et al., 2014, Genome Reference Consortium]. We align non-host reads to the four dengue virus reference genomes (DENV-1, NC 001477; DENV-2, NC 001474; DENV-3, NC 001475; DENV-4, NC 002640) using [BWA-MEM [Li, 2013] / Bowtie 2 [Langmead and Salzberg, 2012]] and independently perform de novo assembly using metaSPAdes [Nurk et al., 2017] to assess potential reference bias. We generate consensus sequences using iVar [Grubaugh et al., 2019] and SAMtools [Danecek et al., 2021], retain positions with a minimum depth of [10*×*], and mask positions with lower coverage as N. We retain only genomes that cover at least 90% of the complete open reading frame for downstream analyses. We assign serotype and genotype using the Dengue Virus Typing Tool in Genome Detective [Vilsker et al., 2019, Fonseca et al., 2019] and confirm the assignments phylogenetically: we use MAFFT v7 [Katoh and Standley, 2013] to align the sequences with reference sequences that we retrieve from GenBank, and we use IQ-TREE v2 [Minh et al., 2020] to reconstruct maximum-likelihood trees under the substitution model that ModelFinder selects [Kalyaanamoorthy et al., 2017], with one thousand ultrafast bootstrap replicates [Hoang et al., 2018].

### 3.2 GISAID sequence data

Our analysis incorporates novel sequences from Ivory Coast (*n* = 24) collected during the period of 2017–2023, alongside all publicly available DENV-1, DENV-2, and DENV-3 sequences retrieved from the GISAID genome database [Khare et al., 2021], accessed on July 26, 2024. The GISAID data summary is provided in Supplementary document S10. During data collection, we apply quality control criteria by excluding sequences with more than 25% ambiguous nucleotides (Ns) or with lengths shorter than 8,250 base pairs (corresponding to less than 75% of the complete dengue virus genome). We also collect associated metadata, including collection date, region, country, location, strain name, serotype, and genotype. We exclude sequences missing essential metadata (such as collection date or country) from further analysis.

We perform multiple sequence alignment using MAFFT version 7 [Katoh and Standley, 2013], integrated via Nextclade v3.10.0 [Aksamentov et al., 2021]. We use reference sequences from the NCBI (National Center for Biotechnology Information) genome database (accession numbers: DENV-1—U88536, DENV-2—U87411, and DENV-3—AY099336) [Sayers et al., 2022] to guide the alignment. We manually review and refine the aligned sequences for quality using AliView version 1.28 [Larsson, 2014]. We then detect recombination using RDP4 [Martin et al., 2015] and remove the identified recombinant regions from the sequences. This results in one DENV-3 sequence that is less than half the full genome length being excluded from the analysis.

To obtain a more balanced dataset, we subsample sequences uniformly across time and geographic locations using SAMPI (https://github.com/jlcherry/SAMPI) [Li et al., 2023], selecting 4, 6, and 8 sequences per year per country for DENV-1, DENV-2, and DENV-3, respectively. We use IQ-TREE v2 [Minh et al., 2020] to infer unrooted maximum-likelihood phylogenetic trees with 1,000 ultrafast bootstrap replicates [Minh et al., 2013, Hoang et al., 2018] under the general-time-reversible (GTR) substitution model. We model among-site rate heterogeneity using a discrete gamma distribution with four categories and an estimable proportion of invariant sites. We then assess the presence of a temporal signal in the sequence data using TempEst v1.5.3 [Rambaut et al., 2016]. Before conducting subsequent phylogenetic analyses, we filter out sequences that exhibited weak temporal structure. The final datasets comprise 1,131 DENV-1 sequences, 1,655 DENV-2 sequences, and 1,152 DENV-3 sequences. Supplementary Figure S1 presents the country-specific numbers of DENV sequences included in the analysis.

### 3.3 Epidemiology of dengue

To complement our phylodynamic analysis, we retrieve national-level dengue case measurements from *Our World in Data* (OWID; [Our World in Data, n.d.a]), accessed in July 2026. The dataset is based on estimates from the Institute for Health Metrics and Evaluation (IHME) Global Burden of Disease (2025), with substantial processing by OWID. It provides annual estimates of new dengue cases per country from 1990 to 2023. We extract case counts for Ivory Coast and neighboring countries to explore epidemiological trends and assess their correspondence with our phylogeographic findings.

### 3.4 Covariate data sources

We collect the number of dengue cases of each country mostly from the World Health Organization’s (WHO) Dengue Global dashboard ([World Health Organization, n.d.b]). For countries where WHO data are unavailable, we first use incidence records from Our World in Data ([Our World in Data, n.d.b]), covering the Democratic Republic of the Congo, Venezuela, Papua New Guinea, the Philippines, Japan, Nigeria, Taiwan, South Korea, Haiti, Brunei, Somalia, Comoros, Djibouti, Nauru, Guam, American Samoa, Bhutan, Egypt, and the United Arab Emirates. When data gaps persist, we rely on additional region-specific resources: a paper for Reunion [Vincent et al., 2023]; Fit for Travel advisories for Saint Barthelemy, Guadeloupe, Saint Martin, and Martinique ([Fit for Travel, n.d.]); WHO situation reports for New Caledonia and the Wallis and Futuna Islands ([World Health Organization, n.d.a]); TravelHealthPro for French Polynesia ( [TravelHealthPro, n.d.]); a systematic review paper for Mayotte [Hafsia et al., 2022]; WHO outbreak alerts for French Guiana ([World Health Organization, n.d.c]); and PAHO surveillance data for Anguilla ([Pan American Health Organization, n.d.]). For the phylogeographic analysis, we transform dengue incidence as log(dengue cases/100,000 + 1). The addition of one accommodates countries with zero reported cases. We apply the log transformation because the effect of dengue incidence is expected to scale with relative rather than absolute changes, yielding a more approximately linear relationship with the response. We present the country-specific dengue case data in Supplementary Figure S2. Supplementary Figure S6 displays the corresponding destination–origin differences in log-transformed dengue case data.

For the annual precipitation of a country, we base the majority of our precipitation dataset on Our World in Data ([Our World in Data, n.d.c]). For countries not included in that dataset, we refer to additional sources. Taiwan’s rainfall data are obtained from its Central Weather Administration ([Central Weather Administration, n.d.]). Data for Singapore are retrieved from the Meteorological Service Singapore ([Meteorological Service Singapore, n.d.]). Additional sources include Trading Economics for Mayotte ([Trading Economics, n.d.a]) and the World Bank Climate Knowledge Portal for Sao Tome and Principe ([World Bank, n.d.a]). Further precipitation data are obtained from Climate.top for French Guiana ([Climate.top, n.d.]), from Climates to Travel for Reunion, Palau, and Wallis and Futuna ([Climates to Travel, n.d.]), and from Weather & Climate for Guadeloupe, Martinique, Guam, Tonga, and the Maldives ([Weather & Climate, n.d.]). Supplementary Figure S8 presents the pairwise destination–origin differences in annual precipitation.

For the country-level temperature covariate, we primarily use monthly average surface temperature data from Our World in Data’s “Monthly average surface temperatures by year” dataset ([Our World in Data, n.d.d]). Because our analysis focuses on seasonal suitability for dengue transmission, we consider using summer temperature instead of annual mean temperature. For countries or territories in the Northern Hemisphere, we calculate summer temperature using the average of June, July, and August, while for those in the Southern Hemisphere, we calculate summer temperature using the average of January, February, and December. For countries or territories not covered in the Our World in Data dataset, we supplement the data using Berkeley Earth regional average temperature records, including Taiwan, Mayotte, Guadeloupe, Reunion, Saint Martin, Barbados, French Guiana, Grenada, Martinique, Saint Kitts and Nevis, Saint Lucia, Guam, Palau, and Tonga ([Berkeley Earth, n.d.]). When Berkeley Earth data are also unavailable or incomplete, we use additional publicly available climate resources, including Singapore’s open data portal ([Government of Singapore, n.d.]), WeatherSpark for Maldives and Wallis and Futuna ([WeatherSpark, n.d.]), and Weather and Climate for Nauru and Saint Barthelemy ([Weather and Climate, n.d.]). We show the pairwise destination–origin differences in country-specific mean summer temperature in Supplementary Figure S7.

We collect absolute latitude data primarily from the Google Developers dataset ([Google Developers, n.d.]). Since latitude is intended to capture geographic variation associated with distance from the equator rather than hemispheric direction, we use its absolute value so that comparable locations in the Northern and Southern Hemispheres are treated equivalently. For countries not fully covered by this source, we use GeoDatos to obtain data for Saint Barthelemy, Saint Martin, Sao Tome and Principe, and Wallis and Futuna ([GeoDatos, n.d.]). We retrieve data for Saint Kitts and Nevis from the Climate Knowledge Portal ([World Bank, n.d.b]), and for additional details on Nepal from Trading Economics ([Trading Economics, n.d.b]).

### 3.5 Bayesian phylodynamic and phylogeographic analysis

We conduct Bayesian timed-tree estimation in BEAST X v1.10.5 [Baele et al., 2025] using an uncorrelated lognormal relaxed molecular clock [Drummond et al., 2006] , and the GTR nucleotide substitution model [Tavaŕe, 1986] with gamma-distributed among-site rate variation [Yang, 1993] for DENV-1, DENV-2, and DENV-3. For the DENV-1 and DENV-2 datasets, we specify a non-parametric Skygrid coalescent prior [Gill et al., 2013] with grid points spaced at two-year intervals. We initially applied the same Skygrid prior to DENV-3, but the resulting MCMC showed poor mixing and failed to achieve satisfactory convergence. We therefore resort to a parametric exponential coalescent prior for DENV-3.

After performing the evolutionary analysis to obtain a posterior distribution of phylogenetic trees, we follow the approach of Pagel et al. [2004] and retain a sample of empirical trees. Then we conduct phylogeographic analysis integrating over those sampled trees. We adapt a Bayesian generalized linear model (GLM) to parameterize the country-to-country transition rates in the discrete phylogeographic model [Lemey et al., 2014]. Specifically, we develop a log linear regression model to regress the infinitesimal transition rate from location *i* to location *j* on the following predictors: log-transformed origin sample size (the number of dengue virus sequences from location *i* included in our analyzed dataset), log-transformed destination sample size (the number of dengue virus sequences from location *j* included in our analyzed dataset), absolute origin latitude, absolute destination latitude, annual precipitation difference, mean summer temperature difference, and log-transformed dengue case/100,000 difference, where each difference variable is defined as the value for destination location *j* minus the value for origin location *i*. In the phylogeographic analysis, we use the transformed covariates log(dengue cases/100,000 + 1) and log(dengue cases/100,000 + 1) to account for the possibility of zero counts in some countries. The corresponding regression coefficients are denoted as β_log(origin sample size)_, *β*_log(destination sample size)_, *β*_absolute origin latitude_, *β*_absolute destination latitude_, *β*_annual precipitation difference_, *β*_summer temperature difference_, and β_log(dengue cases/100,000) difference_.

We perform three independent Markov chain Monte Carlo (MCMC) simulations of 200 million steps each to estimate the Bayesian phylogeographic posteriors, sub-sampling every 20,000 generations and discarding at least the first 10% as burn-in. We assess model convergence and chain mixing using Tracer v1.7.3 [Rambaut et al., 2018]. We combine the log files with LogCombiner v1.10.4 [Suchard et al., 2018], ensuring that all parameters achieved effective sample sizes (ESS) greater than 200. We summarize the phylogeographic tree samples with TreeAnnotator v1.10.4 [Suchard et al., 2018] to obtain a maximum clade credibility (MCC) tree and visualize the resulting MCC tree using FigTree v1.4.4 [Rambaut, 2018].

## 4 Results

### 4.1 Dengue virus phylogeography

We classify the study and background sequences as DENV-1 (*n* = 1131), DENV-2 (*n* = 1655), and DENV-3 (*n* = 1152). The DENV-1 sequences originate from 77 countries (12 from the Ivory Coast), the DENV-2 sequences originate from 75 countries (6 from the Ivory Coast), and the DENV-3 sequences originate from 69 countries (13 from the Ivory Coast). Bayesian phylodynamic analysis of the study and background datasets, summarized in the MCC trees shown in Supplementary Figures S3–S5, shows that the Ivory Coast sequences belong to two distinct DENV-1 clades, one DENV-2 clade, and one DENV-3 clade. The larger DENV-1 cluster includes 132 sequences from 25 countries (11 from the Ivory Coast). The smaller DENV-1 cluster comprises 19 sequences from 9 countries (1 from the Ivory Coast). The DENV-2 dataset includes 34 sequences from 7 countries, while the DENV-3 dataset includes 35 sequences from 5 countries.

To gain insight into the evolutionary history and spatio-temporal dynamics of DENV in the Ivory Coast, we perform Bayesian phylogeographic reconstruction of all serotypes circulating in the country. As shown in Figure 1, we observe six independent DENV-1 introductions into Ivory Coast. For each introduction, we report the posterior probability for the most likely ancestral location (LP), defined as the probability that a specific location is inferred at the corresponding ancestral node, and the posterior probability for monophyly (PP). The first introduction belongs to lineage 1*V II* and originates from India (LP = 0.71; PP *>* 0.99). It occurs on September 23, 1977 (posterior mean), with a 95% highest posterior density interval (HPDI) ranging from December 3, 1968 to November 28, 1986. The second through the fifth introductions fall within lineage 1*III_A_*. The second introduction originates from Burkina Faso (LP *>* 0.99; PP = 0.33), occurring on April 4, 2017 (95% HPDI: August 4, 2016 – November 5, 2017). The third introduction derives from Burkina Faso (LP *>* 0.99; PP *>* 0.99), occurring on May 9, 2017 (95% HPDI: December 19, 2016 – August 28, 2017). The fourth introduction likewise stems from Burkina Faso (LP *>* 0.99; PP = 0.99), occurring on August 12, 2017 (95% HPDI: October 8, 2016 – July 24, 2018). The fifth introduction also arises from Burkina Faso (LP *>* 0.99; PP *>* 0.99), occurring on October 23, 2017 (95% HPDI: February 21, 2017 – June 25, 2018). Finally, the sixth introduction belongs to lineage 1*III_A._*_2_ and originates from Nigeria (LP = 0.47; PP *>* 0.99), occurring on July 24, 2019 (95% HPDI: June 5, 2018 – September 8, 2020).

**Figure 1:**
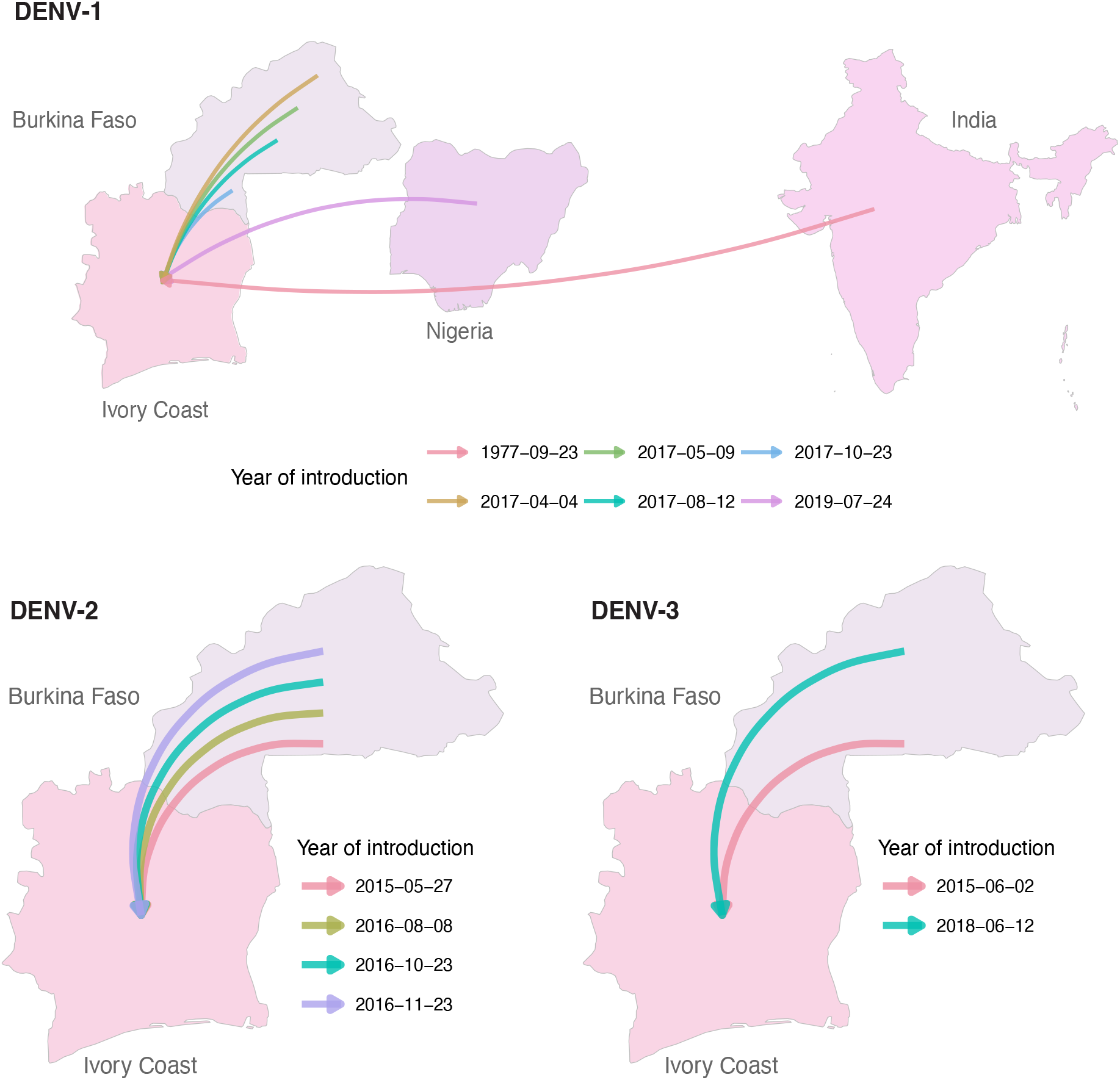
DENV-1, DENV-2, and DENV-3 flux into Ivory Coast.

For DENV-2, Figure 1 demonstrates four independent introductions into Ivory Coast, all originating from Burkina Faso and associated with lineage 2*II_B_*. Each introduction is supported by high location probabilities above 0.85 and mostly strong posterior support (PP = 0.91 for the first, PP = 0.22 for the 2nd introduction, PP *>* 0.99 for the 3rd introduction, and PP = 0.98 for the 4th introduction). The first introduction occurs on May 27, 2015 (95% HPDI: October 22, 2014 - October 22, 2015), the second on August 8, 2016 (95% HPDI: May 25, 2016 - December 19, 2016), the third on October 23, 2016 (95% HPDI: May 21, 2016 - March 27, 2017), and the fourth on November 23, 2016 (95% HPDI: July 14, 2016 - April 2, 2017).

For DENV-3, Figure 1 illustrates two independent introductions into Ivory Coast, both originating from Burkina Faso. The first introduction is associated with lineage 3*III_B_.*2.2, while the second is associated with lineage 3*III_B_.*2.1 These events are all supported by high location probabilities (LP = 0.99) and high posterior probabilities (PP *>* 0.99). The first introduction occurs on June 2, 2015 (95% HPDI: May 6, 2014 - May 18, 2016). The second introduction occurs on June 12, 2018 (95% HPDI: August 20, 2016 - July 27, 2020). We have the estimated Bayesian MCC tree of DENV-1 shown in Figure S3, DENV-2 in Figure S4, and DENV-3 in Figure S5.

### 4.2 Cryptic circulation of dengue virus

Our dataset contains six distinct lineages (Figure 2). Specifically, three lineages belong to DENV-1 (1*III_A_*, 1*III_A._*_2_, and 1*V II*), one lineage belongs to DENV-2 (2*II_B_*), and two lineages belong to DENV-3 (3*III_B_.*2.1 and 3*III_B_.*2.2).

**Figure 2:**
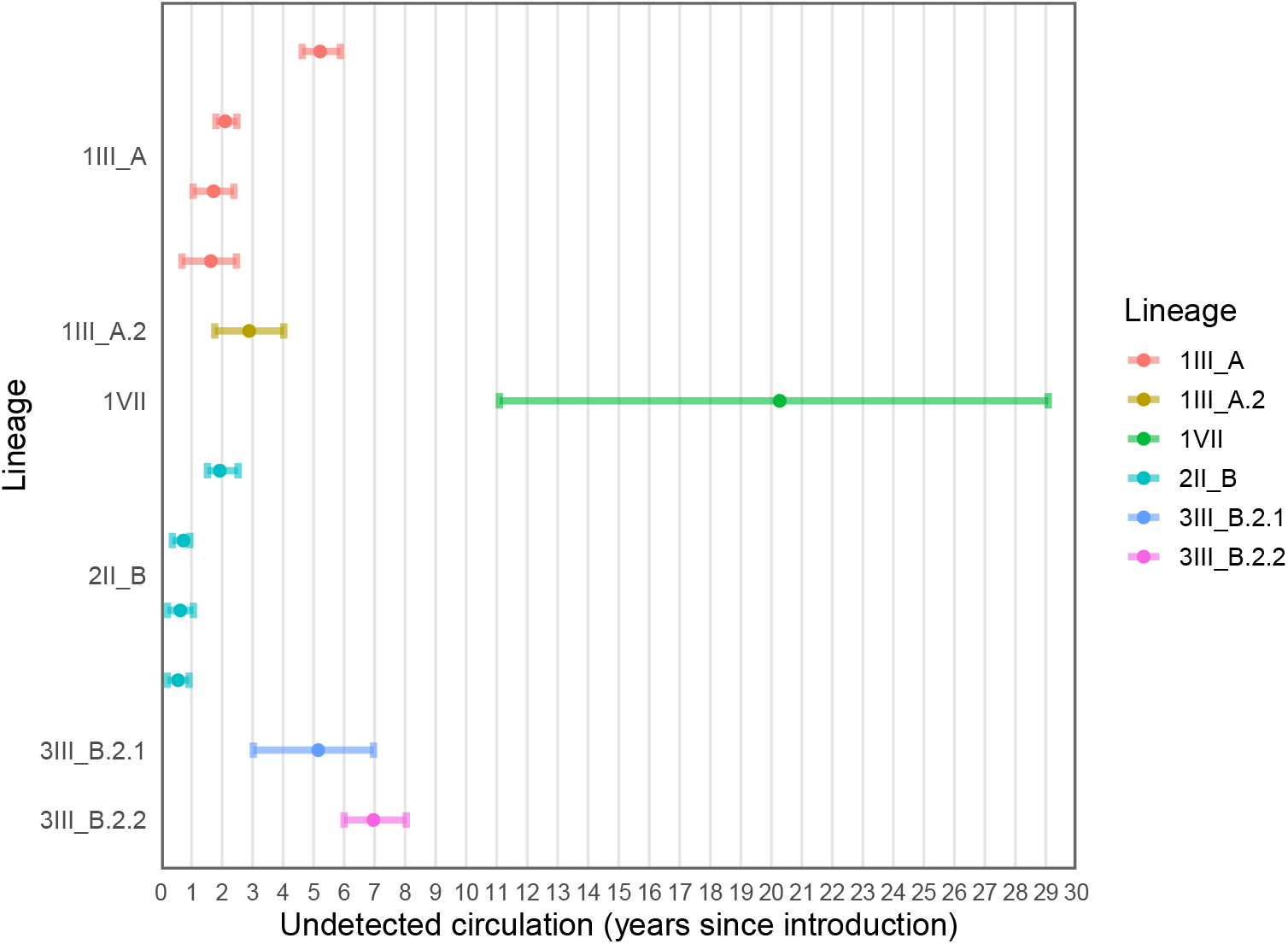
DENV-1, DENV-2, and DENV-3 flux into Ivory Coast.

The introduction of lineage 1*V II* from India into Ivory Coast in 1977 exhibits the greatest uncertainty and the longest period of cryptic circulation, defined as the time interval between the inferred introduction date and the earliest sampling date among sequences descending from that introduction. Overall, DENV-1 shows the greatest lineage diversity and the widest range of cryptic circulation durations among the three serotypes, ranging from 0.7 to 29 years, compared with 0.2–2.5 years for DENV-2 and 3–8 years for DENV-3. In contrast, DENV-2 is represented by a single lineage (2*II_B_*), and the inferred introductions display relatively short periods of cryptic circulation with reduced temporal uncertainty. DENV-3 includes two distinct lineages across two introductions (3*III_B_.*2.1 and 3*III_B_.*2.2), with periods of undetected circulation and associated uncertainties that are larger than those observed for DENV-2 and most DENV-1 introductions.

### 4.3 Predictors of spatial transition rates

Across all three serotypes, the log-transformed origin and destination sample sizes receive the strongest posterior support as predictors of country-to-country transition rates (Figure 3). Both predictors have posterior inclusion probabilities close to 1 for DENV-1, DENV-2, and DENV-3, with consistently positive conditional regression coefficients. For log-transformed origin sample size, the posterior mean coefficients are 0.74 (95% HPDI: [0.56,0.92]), 0.65 (95% HPDI: [0.50,0.79]), and 0.62 (95% HPDI: [0.47,0.76]) for DENV-1, DENV-2, and DENV-3, respectively. The corresponding coefficients for log-transformed destination sample size are 0.86 (95% HPDI: [0.73,0.97]), 0.90 (95% HPDI: [0.78,1.01]), and 0.66 (95% HPDI: [0.52,0.80]) for DENV-1, DENV-2, and DENV-3, respectively.

**Figure 3:**
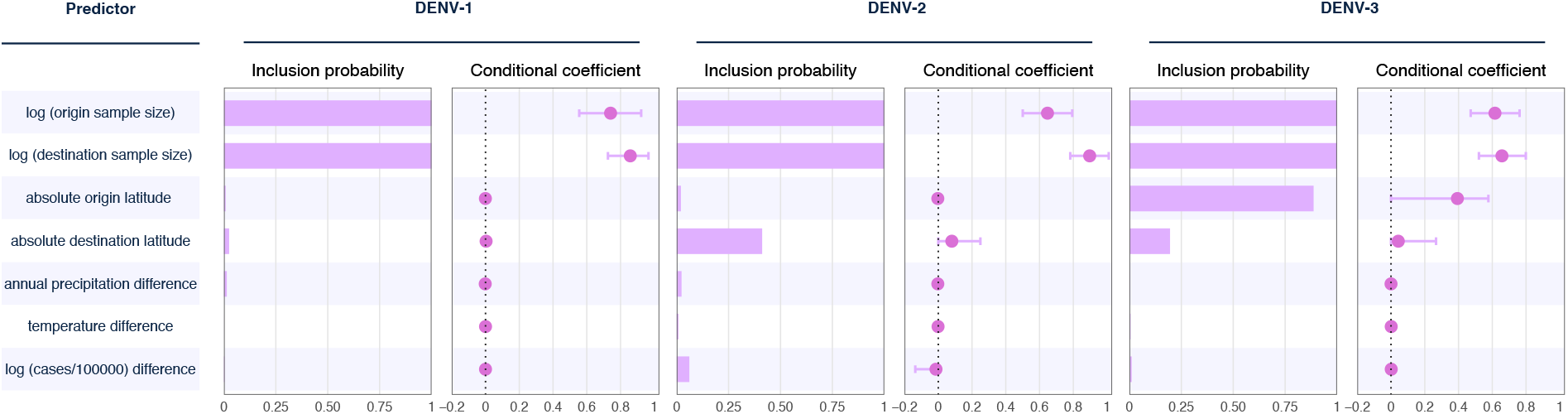
Predictors of dengue fever infections.

Absolute origin latitude shows substantial support as a predictor of DENV-3 transition rates, with a posterior inclusion probability of 0.89 and a positive posterior mean coefficient of 0.39 (95% HPDI: [0,0.58]). In contrast, absolute destination latitude shows weaker support, with posterior inclusion probabilities of 0.41 for DENV-2 and 0.2 for DENV-3. The corresponding posterior mean coefficients are 0.08 (95% HPDI: [0,0.25]) and 0.04 (95% HPDI: [0,0.27]), respectively. Absolute destination latitude shows negligible support for DENV-1.

The environmental and epidemiological difference predictors generally show little support. Annual precipitation difference and mean summer temperature difference have posterior inclusion probabilities close to zero for all three serotypes, and their posterior distributions concentrate near zero. Similarly, the difference in log-transformed dengue cases per 100,000 population shows little evidence of an association with transition rates. DENV-2 shows only weak support for this predictor, with a posterior inclusion probability of 0.06 and a posterior mean coefficient of -0.01 (95% HPDI: [-0.14,0]).

## 5 Discussion and conclusion

Our study reports 24 newly collected dengue virus sequences from Ivory Coast, including 8 DENV-1, 3 DENV-2, and 13 DENV-3 sequences. The DENV-1 and DENV-3 samples are collected between 2022 and 2023, while the DENV-2 sequences date back to 2017. Prior to this study, only 7 dengue virus sequences from Ivory Coast are available on GISAID[Khare et al., 2021]. By adding these 24 novel sequences, we have nearly quintupled the number of publicly available dengue virus genomes from the country. Combining these new sequences with a representative background dataset, our study is based on the most comprehensive dengue virus dataset from the Ivory Coast to date. Supplementary Figure S9 shows the geographic distribution of all 31 Ivory Coast sequences.

Our reconstruction of the viral transmission dynamics reveals that almost all detected dengue virus importation events, across all serotypes, originated from Burkina Faso. This observation is consistent with prior studies Suzuki et al. [2017], which found that a dengue virus strain isolated from a traveler returning to Japan from Ivory Coast in 2017 is homologous to the 2016 strain from Burkina Faso. This finding is epidemiologically plausible given the geographic proximity between the two countries. Burkina Faso and Ivory Coast share a 545-kilometer border, facilitating cross-border transmission of the virus. This finding is also supported by surveillance data.

As illustrated in Figure 4, estimated dengue case counts in Burkina Faso and Ivory Coast exhibit broadly similar temporal patterns, with both countries experiencing an overall increase between 1990 and 2023. This epidemiological pattern is consistent with our Bayesian phylogeographic reconstruction, which identifies Burkina Faso as the predominant source of DENV introductions into Ivory Coast. Of the 12 independent introductions inferred across the three serotypes, 10 originate from Burkina Faso: four of the six DENV-1 introductions and all four DENV-2 and both DENV-3 introductions. Moreover, among the 11 introductions with estimated dates in 2015 or later, 10 are traced to Burkina Faso. Together, these findings indicate that recent DENV circulation into Ivory Coast has been shaped largely by repeated introductions from Burkina Faso across multiple serotypes and viral lineages, rather than by a single isolated transmission event.

**Figure 4:**
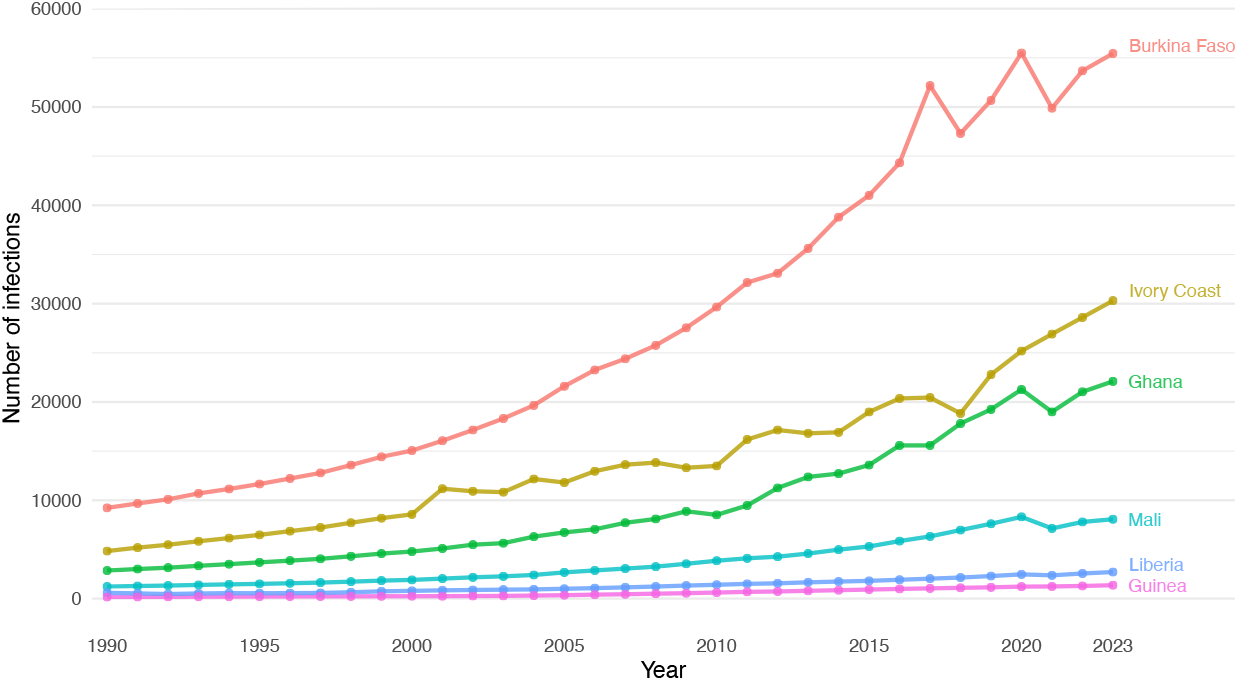
Dengue fever infections in Ivory Coast and neighboring countries.

Among the three locations shown in Figure 4, Burkina Faso has the highest estimated dengue burden in recent years, followed by Ivory Coast, whereas Guinea has substantially fewer cases. By the end of the study period (2023), the estimated number of cases in Burkina Faso is nearly twice that in Ivory Coast and approximately 40 times that in Guiana. However, the availability of GISAID data does not reflect these differences in disease burden as fewer sequences from Burkina Faso than from French Guiana were available for DENV-2, and fewer sequences from Burkina Faso than from Ivory Coast were available for DENV-3.

Despite its relative genomic underrepresentation, Burkina Faso is repeatedly inferred as the ancestral source of nearly all recent introductions into Ivory Coast. This consistency across DENV-1, DENV-2, and DENV-3 strengthens the evidence that Burkina Faso plays a central role in regional DENV circulation. Nevertheless, the true number and diversity of introductions from Burkina Faso most likely remain underestimated.

Furthermore, our analysis suggests that the earliest introduction of the dengue virus into Ivory Coast likely originated from India around 1977. This importation is assigned to lineage 1*V II* and is associated with huge uncertainty. This finding is consistent with the limited availability of sequence data from the late 20th century. The earliest published report we found on dengue introduction to the Ivory Coast dates to 1998 Durand et al. [2000], when DENV-1 was isolated from a French soldier living in Abidjan.However, this particular sequence is not available in genetic databases and therefore could not be included in our analysis. Nevertheless, it is plausible that French soldiers may have facilitated the transmission of dengue virus from India to the Ivory Coast during that period. At the time, France had only recently relinquished its remaining colonial establishments in India, which were transferred in practice in 1954 and legally in 1962 [Government of India and Government of France, 1954, French Republic, 1962], while the Ivory Coast had gained independence from France in 1960 and subsequently maintained close cooperation and defense ties with France [French Republic, 1961]. It is likely that France still maintained a military presence in both regions, creating a possible pathway for viral transmission.

Most of the dengue virus importation events occurred within the past ten years. This does not imply that transmission from Burkina Faso to Ivory Coast only began recently; rather, it reflects the limited availability of recorded viral sequences from Ivory Coast prior to this period. Our analysis also indicates that dengue virus transmission is not confined to neighboring countries. Increasing global connectivity, driven by historical colonial ties and modern trade, plays a significant role in the spread of viruses. While geographic proximity remains a key factor in cross-border transmission, long-distance introductions facilitated by global movement are also critical to understanding the broader dynamics of dengue virus dissemination.

Despite efforts to account for sampling bias, the GLM results suggest that heterogeneity in genomic sampling explains substantially more variation in the reconstructed country-to-country transition rates than the environmental and epidemiological predictors considered here. Transitions are more likely to be between countries that contribute larger numbers of sequences, likely reflecting the influence of heterogeneous sampling on phylogeographic reconstruction[Lemey et al., 2014].

The positive coefficient for absolute origin latitude may reflect the combined effects of transportation connectivity and uneven genomic surveillance. Many higher-latitude countries represented in our datasets have well-developed sequencing systems and serve as major international travel hubs. Viral lineages introduced into these countries from tropical and subtropical endemic regions are therefore more likely to be detected, sampled, and subsequently observed moving to additional locations. At the same time, limited genomic surveillance in many endemic countries may leave intermediate stages of transmission unobserved, increasing the probability that a well-sampled, higher-latitude country is reconstructed as the origin of an observed transition. Thus, absolute latitude may act as a proxy for international connectivity, surveillance capacity, and geographic sampling heterogeneity rather than directly influencing viral movement.

The weakly negative coefficient for the difference in log-transformed dengue cases per 100,000 population is also broadly consistent with this interpretation. Because we define this predictor as the value at the destination minus that at the origin, a negative coefficient implies higher estimated transition rates when the origin has a higher reported dengue burden than the destination. Countries with intense endemic transmission sustain larger pools of infected individuals and may therefore generate more opportunities for infected travelers to introduce viral lineages into lower-incidence countries. The negative coefficient may therefore reflect the movement of DENV from high-burden tropical and subtropical settings into lower-burden but better-observed destinations. However, this predictor receives very weak posterior support for DENV-2.

Our analyses reveal substantial periods of cryptic dengue circulation in Ivory Coast, with estimated durations ranging from 0.2 to 29 years. These findings suggest that viral transmission may have persisted for years before being detected through routine surveillance. Such undetected circulation likely reflects limitations in case recognition, diagnostic capacity, and genomic surveillance, particularly because dengue infections can be asymptomatic or clinically indistinguishable from other febrile illnesses [Dieng et al., 2022b]. Strengthening sustained molecular surveillance and expanding diagnostic and genomic sampling could therefore improve the early detection of newly introduced and locally circulating dengue lineages in the Ivory Coast.

A key limitation of this study lies in the limited availability of dengue virus genomic data from Ivory Coast, particularly prior to the past two decades. While our analysis includes 24 newly sequenced genomes and incorporates all publicly available Ivory Coast data from GISAID (*n* = 7), the overall number of sequences remains relatively small. This constraint significantly restricts our ability to capture the full evolutionary history and transmission dynamics of dengue virus within the country. Moreover, genomic data from neighboring countries available in GISAID are also sparse and unevenly distributed, where Burkina Faso has 107 sequences, while Ghana, Mali, and Guinea have only 10, 12, and 2 sequences, respectively, and Liberia has none (Figure S1). This uneven genomic surveillance coverage further hampers efforts to characterize regional transmission patterns and cross-border viral flow.

Importantly, such imbalance is not limited to this regional context but is evident at the global scale. We present the number of sequences from each country included in our datasets after subsampling in Supplementary Figures S1 and S2. These figures show that a large number of DENV sequences are available from countries such as the United States, Brazil, China, Thailand, Vietnam, and India, whereas relatively few African countries have sufficient sequence data for analysis. Even after subsampling, this imbalance remains substantial and may affect the robustness of our phylogeographic inferences.

The lack of sequences is especially pronounced for the 20th century, during which few genomic records exist. As a result, it is difficult to determine with confidence how and when the dengue virus was first introduced into Ivory Coast, and how it may have circulated or evolved during that period. Although our phylogeographic analysis suggests a possible introduction from India around 1977, the wide 95% HPD interval around this estimate means that these findings should be interpreted with caution.

This scarcity of historical and current genomic data reflects a broader issue of underinvestment in dengue surveillance in Ivory Coast and surrounding regions. Without consistent sequencing and timely sharing of data, it remains challenging to detect transmission pathways, identify sources of new outbreaks, or monitor viral evolution in real time. Expanding sequencing capacity and improving data availability would not only strengthen regional phylogenetic studies but also provide essential information for guiding public health responses.

This study’s findings underscore the importance of both regional proximity and international connectivity in shaping dengue virus dynamics in West Africa. By expanding the genomic data available for Ivory Coast, our study provides a valuable foundation for future outbreak monitoring, cross-border surveillance, and targeted public health responses. Sustained investment in genomic surveillance and regional data sharing is essential to better understand and control dengue transmission in the region.

## 6 Ethics statement

The samples analyzed in this study were collected as part of the routine national surveillance and outbreak response activities mandated to the Institut Pasteur de Ĉote d’Ivoire by the Ministry of Health and the Ministry of Research. Because the investigation was conducted under the public health response to an ongoing outbreak and used only residual diagnostic specimens, it was exempt from formal ethical review, in accordance with the National Institute of Public Hygiene and as confirmed by the Comité National d’Éthique des Sciences de la Vie et de la Santé (CNESVS). No specimens were collected specifically for research purposes, and clinical management of patients was in no way modified by this study. All personal identifiers were removed from the dataset before analysis: samples were assigned anonymous laboratory codes, and the de-identified epidemiological and clinical data were held separately from any information allowing patients to be re-identified. Only aggregated and anonymised data are reported here.

## 7 Author contributions

Conceptualization: X.T., N.S.T., and M.A.S.; methodology: X.T., N.S.T., and M.A.S.; software: X.T. and M.A.S.; investigation: X.T., H.A.K., S.Y., K.E., and A.E.; resources: H.A.K., S.Y., K.E., and A.E.; data curation: X.T.; formal analysis: X.T.; visualization: X.T.; writing—original draft: X.T.; writing—review and editing: X.T., H.A.K., N.S.T., and M.A.S; supervision: N.S.T. and M.A.S.; funding acquisition: M.A.S.

## 8 Acknowledgements

We thank the researchers and laboratories that generated and shared dengue virus genome sequences through the GISAID databases. XT and MAS are supported through US National Institutes of Health grants R01 AI153044 and R01 AI192139. We acknowledge the parallel computing resources through a generous donation from Advanced Micro Devices, Inc.

## 9 Data availability

All study BEAST XML and posterior tree samples are available at https://github.com/suchard-group/dengue_IvoryCoast_data_code.

## Supplementary Materials

**Figure S1.**
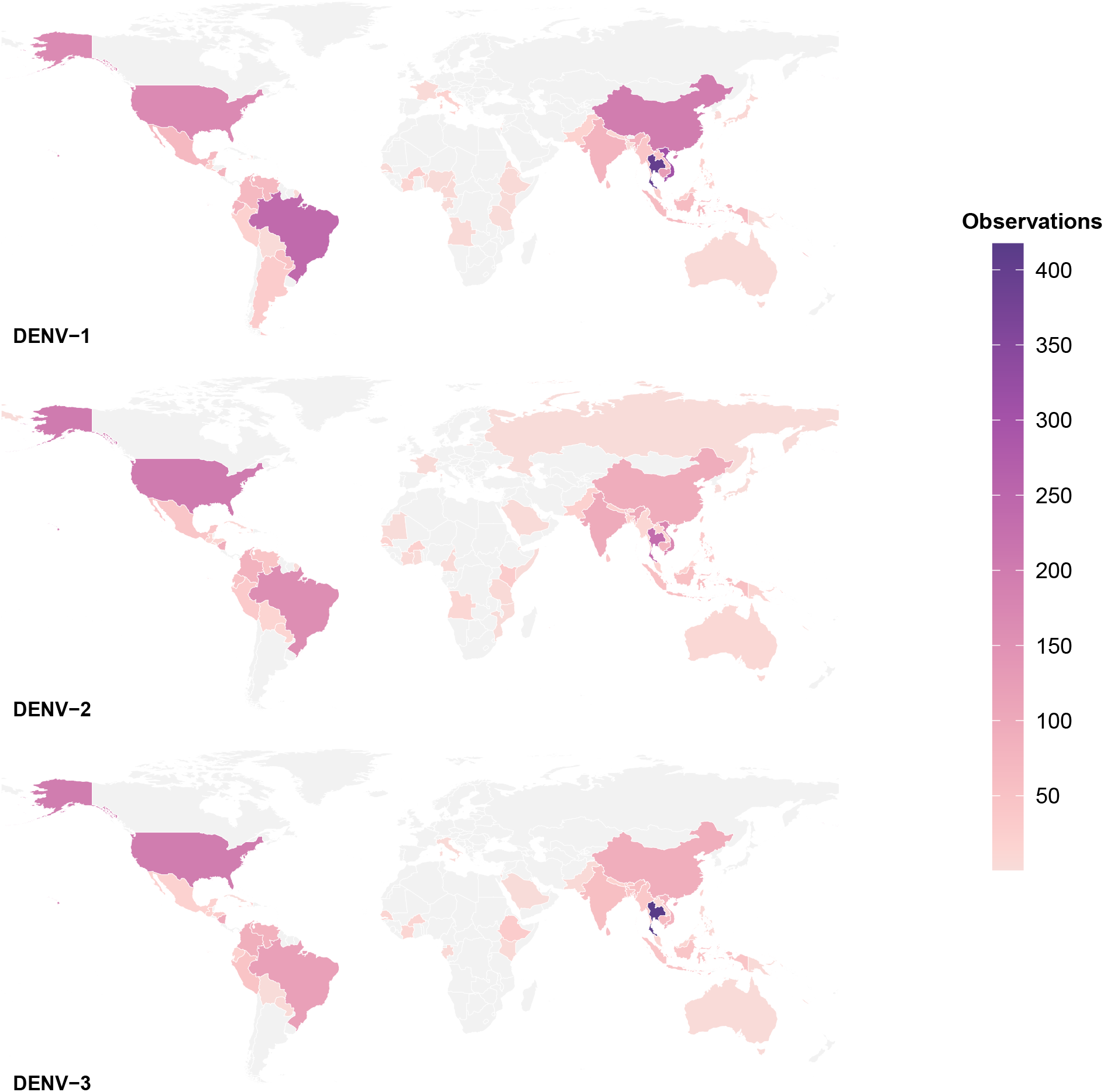
Number of DENV sequences used in the analysis.

**Figure S2.**
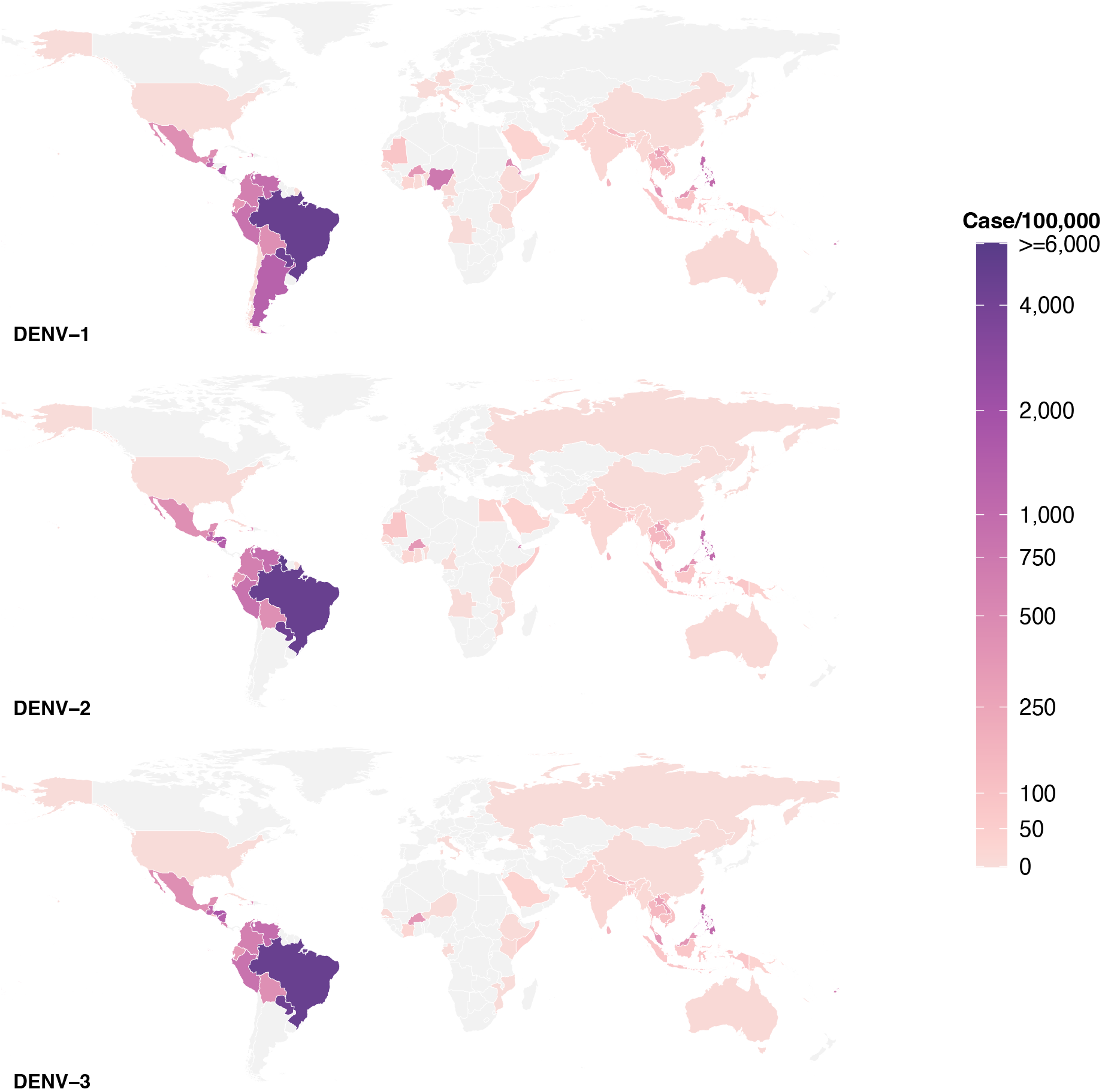
Number of WHO-reported DENV cases/100,000 population.

**Figure S3.**
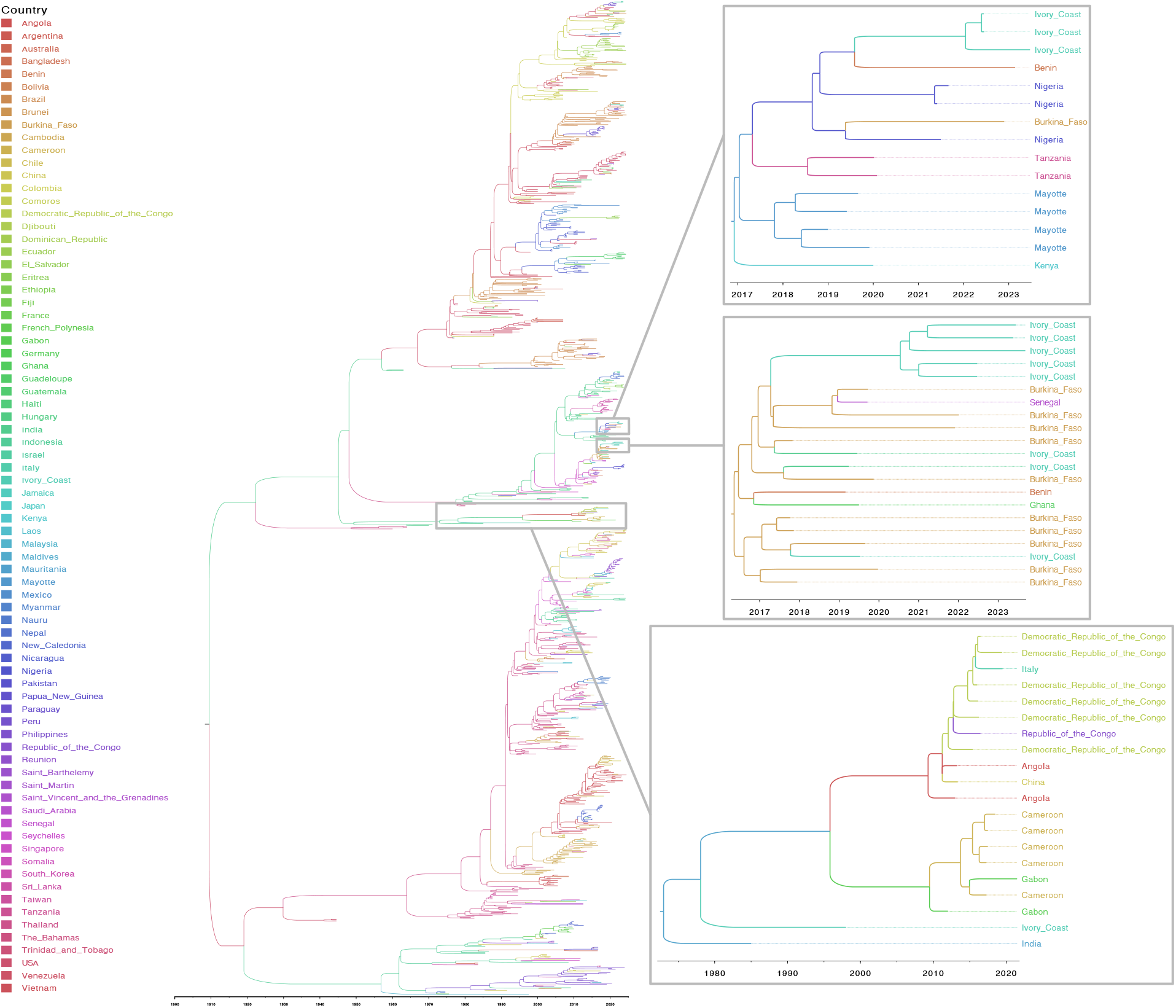
DENV-1 phylogenetic tree and clades including Ivory Coast.

**Figure S4.**
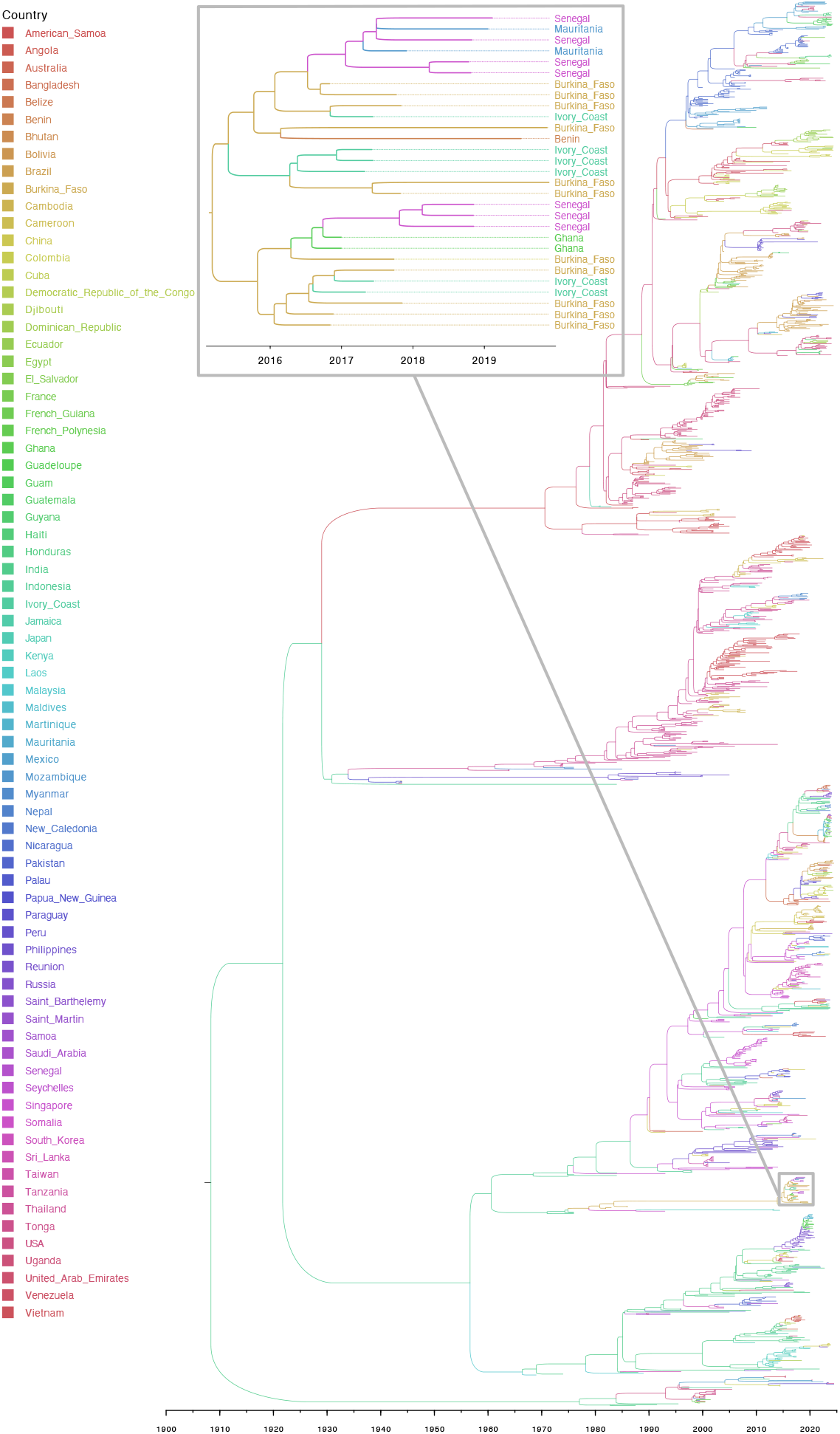
DENV-2 phylogenetic tree and clades including Ivory Coast.

**Figure S5.**
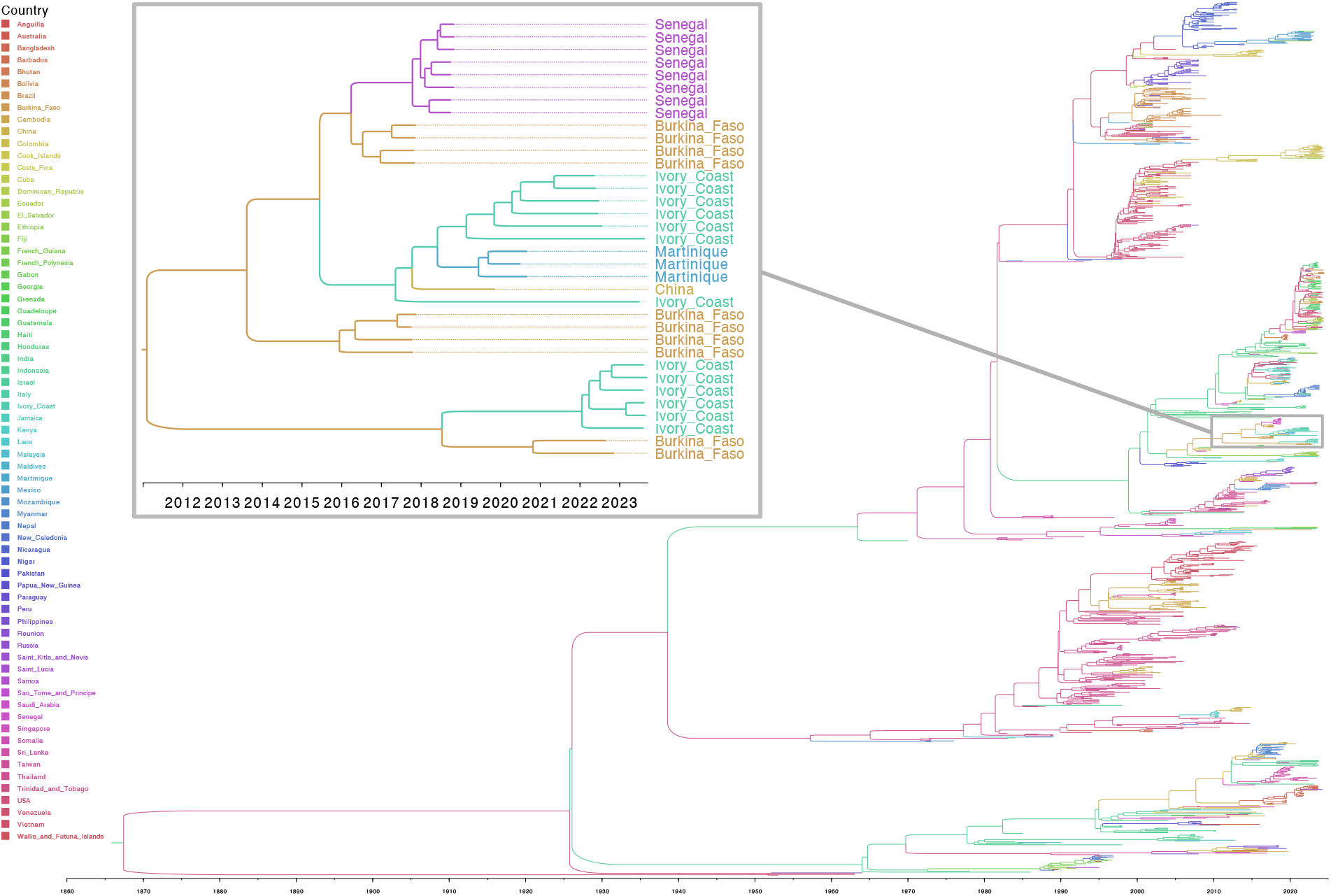
DENV-3 phylogenetic tree and clades including Ivory Coast.

**Figure S6.**
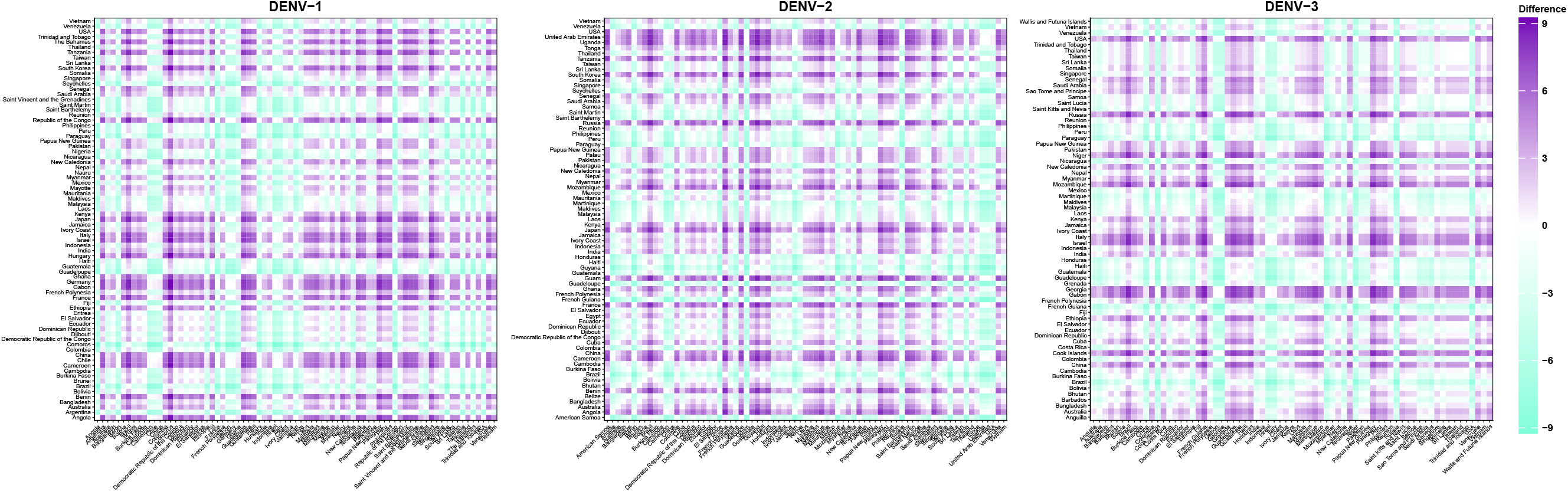
Destination–origin differences in log DENV cases/100,000 population. *Note: Each cell represents the difference between the destination country (column) and the origin country (row)*.

**Figure S7.**
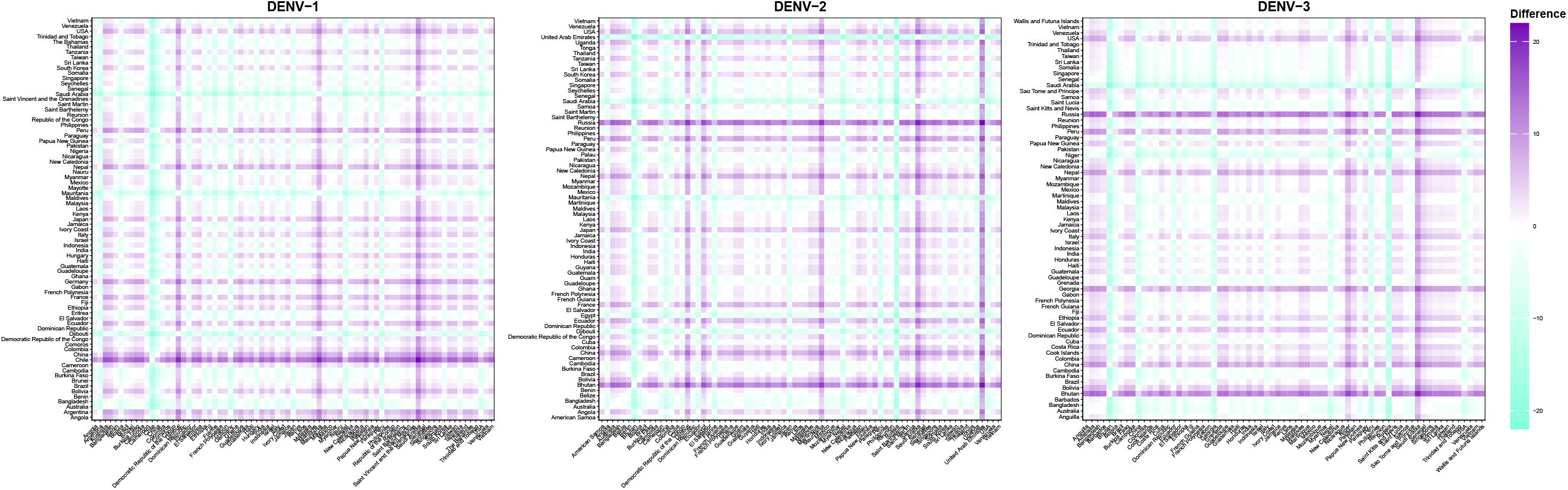
Destination–origin differences in mean summer temperature. *Note: Each cell represents the difference between the destination country (column) and the origin country (row)*.

**Figure S8.**
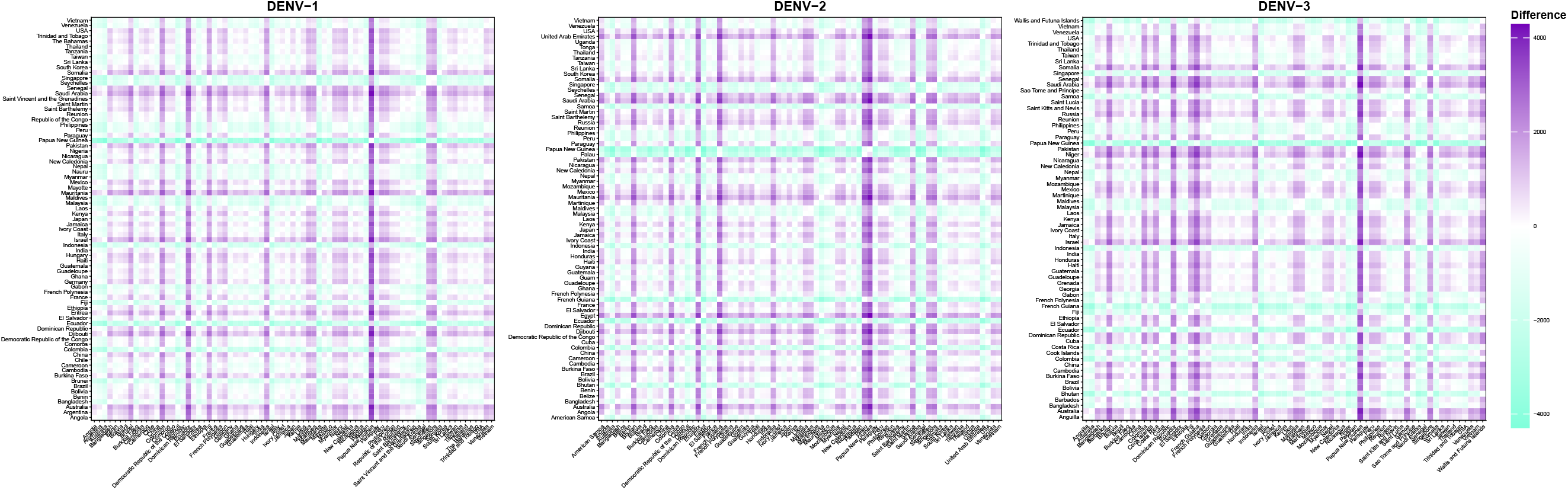
Destination–origin differences in annual precipitation. *Note: Each cell represents the difference between the destination country (column) and the origin country (row)*.

**Figure S9.**
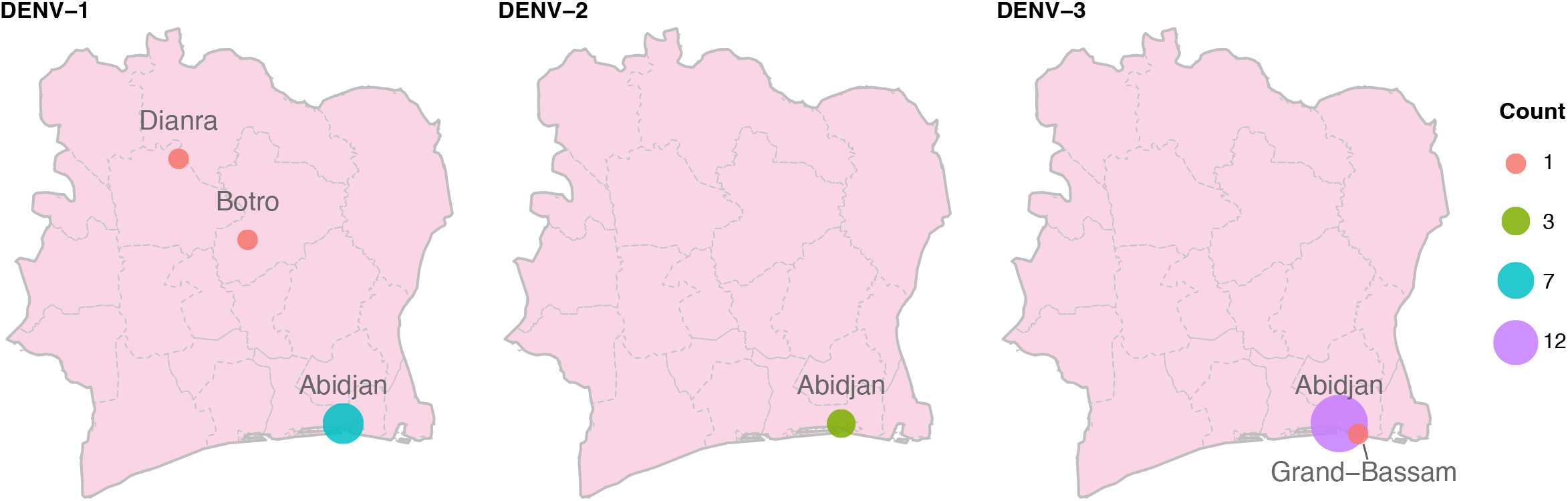
Number of Dengue Virus Sequences Sampled by City in Ivory Coast. *Note: Three DENV-1 sequences and three DENV-2 sequences lack city-level sampling information; only their country of origin, Ivory Coast, is known*.

## Supplementary document S10

### Supplementary Appendix

All genome sequences and associated metadata supporting the findings of this study can be accessed through the persistent digital object identifier https://doi.org/10.55876/gis8.261016gk

In addition to the minted DOI, GISAID also communicates the aggregation of GISAID accession numbers (EPI_ISL_IDs) through the corresponding EPI_SET_261016gk identifier to facilitate both, the acknowledgment of all data contributors and the direct retrieval of the underlying data from GISAID used in this study.

Dengue Virus Data Summary.

| **GISAID Identifier** | **Digital Object Identifier** | **Number of individual viruses** | **Data Collection range** | **Number of countries/territories** |
|---|---|---|---|---|
| EPI_SET_261016gk | https://doi.org/10.55876/gis8.261016gk | 3,899 | 1944-01-01 to 2024-06-26 | 106 |

